# TWO STEPS AWAY FROM EXTINCTION: DO YOU BELIEVE?

**DOI:** 10.64898/2025.12.10.693505

**Authors:** Karina Aparecida Soares de Pádua, Thaís Martins Teixeira, Alison Gonçalves Nazareno

**Author notes:** **CORRESPONDING AUTHORS: KARINA APARECIDA SOARES DE PÁDUA,** Department of Genetics, Ecology, and Evolution - Federal University of Minas Gerais, Avenida Presidente Antônio Carlos 6627, Pampulha, Belo Horizonte, MG, 31270-901, Brazil; **ALISON GONÇALVES NAZARENO,** Department of Genetics, Ecology, and Evolution - Federal University of Minas Gerais, Avenida Presidente Antônio Carlos 6627, Pampulha, Belo Horizonte, MG, 31270-901, Brazil.

## Abstract

Forest fragmentation can have detrimental effects on plant populations, reducing population sizes and depleting genetic diversity, as a consequence. As a matter of urgency, it is crucial to assess the effects of forest fragmentation on genetics and ecological processes, particularly for threatened species on the brink of extinction. Here, we examined the responses of *Dinizia jueirana-facao* G. P. Lewis & G. S. Siqueira (Fabaceae, Caesalpinioideae) – a rare and critically endangered tree species endemic to the Brazilian Atlantic Forest–, to forest fragmentation. Based on theoretical predictions for barochory plant species with small population sizes, we hypothesized that forest fragmentation would reduce gene flow, erode genetic diversity, and negatively impact demography. Using neutral SNPs (Single Nucleotide Polymorphisms) derived from ddRADSeq (double-digest restriction site-associated DNA sequencing), we found that pollen dispersal occurred within short distances, with the majority of outcrossed pollination events occurring locally. Furthermore, contemporary estimates of gene dispersal distances were lower than historical ones, indicating a seasonal shift in the scale of gene flow due to recent forest fragmentation. Our results also indicated no evidence of inbreeding and loss of genetic diversity. In terms of ecological process, the demographic structure of fragmented populations of *D. jueirana-facao* followed a reverse J-shaped size class distribution, with more than 45% of plants found in small diameter classes. While a more in-depth understanding of ecological and evolutionary processes at fine-scale is still needed to safeguard this unique plant species, our infant study plays a crucial role to help keep the evolutionary potential of *D. jueirana-facao*. We stress that the approach used here would be useful to guide conservation and management efforts for species on the brink of extinction.

## INTRODUCTION

Continental-scale analyses of tropical forest fragmentation reveal a predictable and alarming scenario, with an ongoing increase of small and isolated fragments^1,2^. This trend has also been observed in the Brazilian Atlantic Forest, where only 24% of the original vegetation remains, with many forest patches smaller than 50 ha^3–5^. This biome has been historically degraded and deforested by anthropogenic activities such as agricultural expansion, urbanization, and industrialization^6,7^, resulting in highly fragmented landscapes and threatening the long-term persistence of native species as a consequence.

As a matter of fact, habitat fragmentation mainly caused by environmental changes and anthropogenic pressures may have a series of genetic and ecological consequences. For instance, despite being diverse in terms of approaches and contexts, a plethora of studies pointed out the impacts of habitat fragmentation in the loss of standing genetic variation due to genetic drift, limited gene flow, disruption of ecological processes^8–23^. When considering small and isolated plant populations with limited ability to disperse their genes, the negative genetic effects can be even more critical, increasing the amount of correlated crosses, exacerbating the restriction of gene dispersal, intensifying genetic drift, and depleting genetic variation^9,15,18,24–29^. In this context, bringing to attention the needs for effective conservation actions are paramount, especially for critically endangered species on the verge of extinction such as *Dinizia jueirana-facao*, a rare and endemic tree species that occurs only in two small remnant areas of Brazilian Atlantic Forest. To investigate how a critically endangered tree species responds to forest fragmentation, we adopted two approaches. The first one relies on population genomic data obtained from a reduced representation genome sequencing method to assess genetic variation, estimate gene flow, and depict the mating system of *D. jueirana-facao.* The second approach was based on ecological data to investigate demographic structure in *D. jueirana-facao* populations.

Specifically, this methodological framework allowed us to assess the impact of forest fragmentation on genetic diversity, investigate contemporary and historical gene dispersal and mating system patterns, and to test whether *D*. *jueirana-facao* populations are genetically isolated. It also allowed us to compare contemporary and historical estimates of gene flow by testing the hypothesis of genetic diversity loss due to limited gene flow, as predicted to small populations of plant species with limited seed dispersion^9,29^. Additionally, we examine the hypothesis of recent reduction in gene flow scale, as reflected in dispersal distance. This pattern is expected due to the perturbation effect, which is primarily expressed in contemporary estimates, while historical estimates exhibit temporal inertia over a few generations^30,31^. In addition to the genetic component, analysis of demographic size population structure (i.e., diameter size class distribution) may provide information on ecological consequences of forest fragmentation. Accordingly, we assessed whether *D. jueirana-facao* populations fit the reverse J-shaped size class distribution pattern, as it reflects stable recruitment in undisturbed forests, whereas deviations may indicate demographic instability due to fragmentation^32,33^.

While a more in-depth understanding of ecological and evolutionary processes at fine-scale is still needed to safeguard this unique plant species, our infant study plays a crucial role to help keep the evolutionary potential of *D. jueirana-facao*. We stress that the approach used here would be useful to guide conservation and management efforts for species on the brink of extinction.

## MATERIALS AND METHODS

### Studied area

The region where *D. jueirana-facao* occurs has a flat to slightly sloping relief, which makes it attractive for agriculture and characterizes the surroundings of these forests with a sudden transition of landscape with pastures and farmland in addition to the presence of roads. The area of occurrence of the species, which has a history of intense anthropogenic activities dating back 1916, is nowadays designated as a protected area^34,35^.

With only two known localities confined to the Atlantic Forest, the occurrence of *D. jueirana-facao* is limited to southeast Brazil, specifically within the Reserva Natural Vale in the Linhares municipality and on a private property located 12.0 km away from the reserve in the Sooretama municipality, both situated in the Espírito Santo state (Figure S1). The Reserva Natural Vale (RNV) has previously experienced challenges such as illegal collection of plant and animal specimens, predatory hunting, forest fires, and periods of drought^34^. Although the RNV shares borders with other reserves, such as the Reserva Biológica de Sooretama (RBS) and two other private reserves of natural heritage, forming a potential gene flow matrix, its boundaries are surrounded by contrasting environments such as pastures, farmland, and roads.

### The focal plant species

Until 2017, *Dinizia* was considered a monotypic genus, represented by *D. excelsa* Ducke. However, with the description by Lewis et al.^36^, *Dinizia* now comprises two members, including *D. jueirana-facao*, a rare and endemic tree species. Despite being recently described, this tree species is listed as critically endangered^37,38^. It exhibits an emergent canopy ranging from 19 to 40 m and a trunk height of 10 to 22 m, while its congener Amazon tree species can reach heights of more than 60 m. *Dinizia jueirana-facao* features bright yellow inflorescence, hermaphrodite flowers measuring 8.5 – 10 mm, with occasional instances of functionally male flowers preceding the suppression of gynoecium development. On the other hand, *D. excelsa* has whitish green to greenish yellow inflorescences, the flowers are mostly functionally male due to the suppression or absence of the gynoecium. In *D. jueirana-facao* the dehiscent woody fruits are dark brown to black when ripe, measuring 40 – 46 × 8.5 – 10 cm and containing 13 – 15 black seeds that are hard and measure 25 – 30 × 16 – 19 mm. Meanwhile, *D. excelsa* has a wine-red, almost winged, indehiscent fruit, that measures 20.5 – 35 × 4.5 - 8.5 cm, containing 7 – 12 seeds, (10 –)14 – 15 × 6 – 7 mm^36^.

While there is a morphological characterization for *D. jueirana-facao*’s reproductive structures, no information regarding pollination syndrome is currently available for it. However, pollen monads in *D. jueirana-facao* indicate that this tree species may be wind-dispersed^36^. Bees and small beetles emerge as potential pollinators, as they have been reported as pollen dispersers in *D. excelsa* which presents similar floral traits^36,39^.

Based on our field observations, seeds of *D. jueirana-facao* seem to be primarily dispersed by gravity (i.e., barochory).

### Sampling for genetic and ecological analyses

Taking into account the rarity and the limited number of populations, we adopted an ontogenetic methodological framework to evaluate the genetic consequences of forest fragmentation. This approach, commonly applied in conservation genetics for long-lived species in highly modified landscapes, provides valuable insights into how contemporary processes affect genetic diversity and gene flow of small populations. Reproductive adults, intermediate (i.e., juveniles), and seedlings were collected in RNV (116 individuals) and RBS (95 individuals). Leaf samples from 211 individuals were collected and stored in -20 °C until DNA extraction, and the geographic coordinates of each individual were recorded using a Garmin portable GPS (∼30 cm error). We measured the stem circumference of each tree at approximately 1.3 m from the ground using a tape measure. This measurement was then used to calculate the diameter at breast height (DBH) using the formula 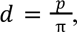 where *p* represents the circumference (the distance around the stem)^40^.

In order to prevent bias and attend the assumptions of each of the performed analyses, *D. jueirana-facao* individuals were categorized into three ontogenetic stages based on DBH: seedlings (DBH **<** 5 cm), intermediate individuals (5 cm > DBH > 14 cm) reproductive adults (DBH ≥ 14 cm) (Table S1). The DBH threshold to classify a tree as reproductive was determined by the lowest DBH observed among all reproductive trees in both populations. Reproductive events were assessed during the field expedition, and additional events were inferred based on the presence of dried reproductive pods and/or seeds under tree canopy.

To distinguish recent from historical genetic patterns and incorporate ecological data, we structured our analyses using three datasets: (a) paternity, spatial genetic structure (SGS), and mating system; (b) genetic diversity and inbreeding; and (c) effective population size (Ne) and population demographic structure. Ontogenetic stage inclusion varied according to the assumptions and hypotheses addressed by each analysis. Full details are provided in Supplementary Table S2.

### DNA extraction, genome library preparation, and sequencing

For each sampled *D. jueirana-facao* individual, total genomic DNA was extracted from leaves using the Macherey-Nagel kit (Macherey-Nagel GmbH & Co. KG), following the manufacturer’s instructions. DNA was quantified using the Qubit dsDNA Assay Kit (Invitrogen). Genomic library preparation for ddRADseq method was performed according to Peterson et al.^41^, with slight modifications to minimize variation of reads among individuals^42^. Digestion reactions were carried out using both *EcoRI* and *MseI* restriction endonucleases (New England Biolabs) on each sample containing 0.5 µg of genomic DNA. After digestion, reactions were purified using the Agencourt AMPure XP system (Beckman Coulter) and eluted in 40 µL water. Subsequently, for each sample, adapter ligation reaction was performed by combining 80 ng of clean digested DNA, 0.35 µM of *MseI* adaptor (common for all samples), 0.50 µM of a sample-specific *EcoRI* double-strand adaptor, 1U of T4 DNA ligase (New England Biolabs), and 1.5 µL of T4 ligase buffer. It was performed in a Bio-Rad thermal cycler at 23 °C for 30 minutes, heated at 65 °C for 10 minutes, and slowly cooled to 23 °C.

The ligation products were purified using the Agencourt AMPure XP system and amplified following the PCR protocol as in Nazareno et al.^42^ Amplicons were pooled and the Pippin Prep system (Sage Science, Beverly, MA) was used to automatically select fragments ranging from 375 to 475 base pairs (bp). Four genomic libraries (100 bp single- end) were sequenced on Illumina HiSeq 2500 (Illumina Inc., San Diego, CA) at The Centre for Applied Genomics in Toronto, Canada.

### SNPs identification and data filtering

Data quality was performed using FastQC program, version 0.11.8^43^. The raw sequence reads files were analyzed using Stacks 2.41 version^44–46^ using *de novo* assembly. In Stacks, process_radtags program was utilized to demultiplex the raw data. This process allowed a maximum of two mismatches in the redemption of barcodes (option -barcode-dist-1 2). Low quality reads and those lacking the expected *EcoRI* site were excluded, and all filtered reads were trimmed to 85 bp. Samples with less than 500,000 reads were excluded from further analyses. Ustacks was employed to align short-read sequences from the output of process_radtags and generate “stacks” (or putative alleles) using a maximum-likelihood framework^47^. The stacks were built with a minimum depth of coverage of three sequences, a maximum number of stacks at a single *de novo* locus, and a maximum distance of two nucleotides between stacks. The -d and -r parameters in Ustacks were used to address over-merged tags and remove highly repetitive stacks, respectively. SNPs were identified using an error rate (option bounded --bound_high 0.1), and the alpha value for the SNP model was set up at 0.05.

Cstacks was utilized to create a catalog, or set of consensus loci, containing all loci from all individuals with three mismatches allowed between loci. The SStacks program was then employed to search the set of stacks in the catalog. In the locus clustering stage, tsv2bam in Stacks was used to establish a data orientation by locus rather than by sample, storing them in standard BAM files. The next step involved using the tsv2Bam outputs in Gstacks to build contigs with single reads, merge the contig with the single-end locus, and align reads from individual samples to the locus. SNPs were identified for each locus and individual in Gstacks, and genotyped SNPs were converted into a haplotype dataset. Finally, the Stacks’s POPULATIONS program^44–46^ was executed to filter and export the SNP dataset to GENEPOP format. Population statistics were computed using the exported SNP dataset. The FASTA files containing per-locus consensus sequences and individual loci sequences were also exported for further analyses.

### Prior genomic analyses

The effect of missing data (MD), mirrored by the number of SNPs, on genetic diversity parameters was examined on Stacks’s POPULATIONS program^44–46^ as recommended by Nazareno and Knowles^31^. The program was running several times varying the MD from 0 to 30% with a fixed minor allele frequency (MAF) equal to 0.004. The effect of MAF, varying from 0.05 to 0.5, on the number of SNPs was also tested. These analyses indicated that 10% of MD is the best threshold to unbiased the results^31^.

Hardy-Weinberg equilibrium (HW) and linkage disequilibrium (LD) tests were performed in R package Pegas version 1.1^48^ and in the Genepop v 4.7.5 software^49^, respectively. Type I error rates for these tests were corrected for multiple comparisons using the sequential Bonferroni procedure^50^. In addition, two filtering analyses were performed. The first one was conducted to exclude non-nuclear loci, aligning per-locus consensus sequences against chloroplast and mitochondrial reference genomes of closely related plant species: *Haematoxylum brasiletto* (NCBI accession number NC_026679.1; NC_045040.1; MN709823.1), *Arapatiella psilophylla* (NCBI accession number MN709845.1), and *Schizolobium parahyba* (NCBI accession number MN709795.1). BLASTn program^51^ was used to identify loci with identity greater than or equal to 80%. Second, the software Bayescan version 2.0^52^ was used to identify locus potentially under selection. Default settings were kept, including 20 pilot runs of 10,000 iterations, a burn-in of 50,000 iterations, and a final run of 100,000 iterations. The prior odds of the neutral model were set up to 10,000 in order to minimize false positives^52^.

### Analyses of parentage, spatial genetic structure, and mating system

The software COLONY version 2.0.6.8^53^ was utilized for the analysis of paternity in cases where there is known maternal kinship but unknown paternal kinship. The dataset was divided into three classes: Seedling Sample, CFS (Candidate Father Sample), and CMS (Candidate Mother Sample) based on the aforementioned ontogenetic stages. All adult trees that passed the raw data quality filters were considered as CFS and/or CMS and the known maternal kinship option in Colony was added as putative mothers.

Putative mothers were determined by analyzing distance matrices between reproductive adults and seedlings based on their geographic coordinates. It was assumed that the putative mother is the breeding tree closest to the progeny, as trees farther away are more likely to contribute to pollen dispersal distance^31,54^. Paternity reconstruction was conducted separately for seedlings from RNV (n=21) and RBS (n=29), using CFS and CMS from both locations (n=85).

The software COLONY was utilized with the following configurations: polygamy for both males and females, inbreeding mating, no clones, monoecious, full likelihood estimates, and medium likelihood precision. The Run Specifications and Sibship Prior parameters were left at their default values. Initially, mistyping error rates (genotyping errors including mutations) and allelic dropout rates were set to 0.001 and 0, respectively. COLONY then re-estimated these rates for each marker locus based on the best reconstructed pedigree.

To calculate seed and pollen dispersal distances, the following criteria were applied: (i) only parents with a paternity or maternity attribution probability greater than 95% were considered, (ii) seedlings with only one putative parent assigned were assumed to be the mother, and this parent was used to infer the seed dispersal distance assuming that the putative mother is the adult reproductive tree closest to the seedling, since those more distant tend to contribute with pollen dispersal distance^31,54,55^, and (iii) for seedlings with two assigned parents, the closest candidate to the seedling was considered the maternal parent, while the other candidate was the paternal parent^54,55^.

Using the obtained paternity results, the direct gene flow (*σ²_rt_*) was calculated using the equation *σ²_rt_ = ½ σ²_p-rt_ + σ²_s-rt_*, where *σ²_p-rt_* represents the pollen dispersal variance and *σ²_s-rt_* represents the seed dispersal variance^56^. Seed (*m_s_*) and pollen (*m_p_*) immigration rates were calculated as the percentage of genotypes not assigned to a candidate parent within the population (with probability > 0.95). In this case, *m_s_* represents the percentage of seedlings without any assigned parent in relation to the total, while *m_p_* represents seedlings with only one assigned parent^57–59^.

### Fine-scale Spatial Genetic Structure

The spatial genetic structure was evaluated using the SPAGeDi program v. 1.5d^60^ through spatial autocorrelation analysis between kinship coefficients and distance intervals among individuals and the *Sp* statistic. Relationship coefficients (*R_ij_*) were computed for each ontogenetic stage of the two localities. The kinship coefficient estimators *F_L_* and six distance intervals were employed, with the jackknife method across all the loci used to calculate the 95% confidence interval (CI) of the standard error of the relatedness coefficients^61^.

The intensity of SGS was assessed using the *Sp*-statistic defined by Vekemans and Hardy^60^. The *Sp*-statistic was used to compare the extent of SGS between localities. The equation *Sp* = − *b* /(1 − *F*_1_), was used to obtain the *Sp*-statistic, where *b* is the regression slope of *F_ij_* on log spatial distance, and *F*_1_ is the mean *F_ij_* between individuals for the first distance class^60^. Significance of the regression was determined by 10,000 permutations of multilocus genotypes. The observed *R_ij_*was considered statistically significant if it deviated from the 95% confidence interval established by data permutations for the null hypothesis of no spatial genetic structure, *R_ij_*=0. Significant positive or negative structure was inferred if the CIs of standard error did not overlap^62,63^.

The gene dispersal distance will also be estimated to compare with the direct gene flow obtained by paternity analysis^56,60^. The root-mean-squared dispersal distance (*σ*) was calculated following the formula of Vekemans and Hardy^60^: *σ*² = *N_b_*/4 *π D_e_*, where the Wright’s neighborhood size (*N_b_*) equals *1/Sp*, and effective population density (*D_e_*) equals product of the census density *D* and *N*_e_*/N*. To calculate *D* we use *D=N/area*, where *N* represents the total number of sampled plants of each locality of *D. jueirana-facao* (RNV= 116, RBS= 95) divided by their distribution area (RNV= 42.99 ha, RBS= 64.81 ha) reported by Lewis et al.^36^ After, to calculate the values of effective population size (*N*_e_) we use the number of reproductive trees sampled in RNV (n=17) and RBS (n=13).

### Mating system analysis

The kinship coefficient, as described by Loiselle et al.^61^ and implemented in the SPAGeDi program^64^, was utilized to estimate random outcrossing rates (1- *s*, where *s* represents the selfing rate). These rates were determined based on standardized identity disequilibrium for adult individuals from both localities of *D*. *jueirana-facao*. Significance for the identity disequilibrium coefficient was obtained through 1,000 permutations, and a jackknife over loci was employed to calculate the standard error of outcrossing estimates.

### Genetic diversity and fixation index parameters

To evaluate the potential impact of forest fragmentation on the genetic variation of *D. jueirana-facao* over distinct generations, we employed a rarefaction method implemented in ADZE 1.0^65^ to estimate standardized allelic richness (*A*) and standardized private allelic richness (*R*). This method accounted for differences in sample sizes between the compared datasets (i.e., reproductive adults before fragmentation and seedlings) of *D. jueirana-facao* in both localities. ADZE was run using a maximum standardized sample size of 29, with a missing data tolerance of 0.15 per locus. In addition, genetic estimates of *N*_e_ for each sampling location were obtained using the linkage disequilibrium method in the program NeEstimator Version 2.166. To include all alleles in the *N*_e_ calculation, we used an allele frequency critical value equal nil^66^. The 95% CI were calculated using the jackknife resampling method.

To represent individuals before (≥120 years old) and after the period of forest fragmentation, we selected reproductive adults and seedlings from RNV (n_adults_=12 and n_seedlings_=21) and RBS (n_adults_=13 and n_seedlings_= 29), based on tree age estimates and historical reports of anthropogenic activities in the study region^34,35^. Tree ages were estimated using DBH data and a growth rate of 0.36 cm.year^-1^ SE±0.19 (DBH > 20 cm) and 0.62 cm.year^-1^ SE±0.19 (DBH < 20 cm). These estimates were based on the growth rate of the congener species *D. excelsa*, which is 0.53 cm.year^-1^ (DBH > 20 cm) and 0.91 cm.year^-1^ (DBH < 20 cm)^67^. Considering that tree species in the Amazon biome, such as *D. excelsa*, exhibit higher growth rates compared to those in the Atlantic Forest^68^, we used a conservative growth rate 32% lower than *D. excelsa*.

To assess the level of inbreeding for the two generations in both localities, we calculated the inbreeding coefficient (Wright’s Fixation Index *F_IS_*)^69^ using the BasicStats function in the R package DiveRsity^70^.

### Population demographic size structure

In order to test whether *D. jueirana-facao* populations fit to a reverse J-shaped size class distribution, we utilized a dataset containing information on the distribution of DBH and tree height from all sampled individuals. First, we tested the normality of these data, applying the Shapiro-Wilk Normality test. After that, we used the Kolmogorov-Smirnov test to verify if there are differences in demographic size structure between ontogenetic stages and localities. Diametric distribution and height graphs were plotted to examine the pattern of demographic size structure. All these statistical analyses were performed in R^71^.

Additionally, densities of seedlings, intermediates, and reproductive adults were used to determine the regeneration status within each locality. The regeneration status was categorized as: (i) good = if seedlings > or < intermediates > reproductive adults; or (ii) fair = if seedlings > or ≤ intermediates ≤ reproductive adults^72^.

## RESULTS

### Sequencing, SNPs identification, data filtering, and prior genetic analyses

Illumina sequencing yielded approximately 144 million single-end raw reads per lane. The mean number of retained reads that met the default quality filters, including a Phred quality score > 33, were as follows: 2,319,619.1 ± 1,076,166.86 SE for the first lane, 2,099,300.824 ± 1,299,075.296 for the second lane, 2,030,593.393 ± 1,702,835.731 for the third lane, and 2,026,080.63 ± 1,357,852.272 for the fourth lane. A total of 12% of individuals (n=27) were excluded from the dataset for having less than 500,000 retained reads (Figure S2; Table S1).

Considerable variation in the number of SNPs was observed for both localities: RNV (MD_0%_=450 to MD_30%_=9,869) and RBS (MD_0%_=506 to MD_30%_=9,481). Genetic diversity parameters showed no sensitivity to MD variation, with overlapping 95% confidence intervals (CI) for each ontogenetic stage (see Table S5). To test the effect of MAF on the number of polymorphic sites, a fixed value of 10% MD was used, resulting in approximately 4,300 SNPs. The number of SNPs due to MAF variation ranged from 11 SNPs (MAF=0.5) to 4,194 SNPs (MAF=0.05) (Table S6).

Prior genomic analyses were performed to eliminate possible bias. For the paternity, SGS, and mating system analyses, an initial dataset consisting of 702 SNPs was identified. After Bonferroni adjustment, 44 SNPs deviated from H-W equilibrium (*p* > 7.12 x 10^-5^) and no SPN showed Linkage Disequilibrium (*k* = 2.46 x 10^5^, *p* < 2.03 x 10^-7^). A total of 14 potential loci under selection were identified and no plastid locus was observed, resulting in a dataset with 644 SNPs. For genetic diversity and inbreeding estimates, a total of 4,738 SNPs were identified. However, 166 SNPs deviated from H-W equilibrium after Bonferroni adjustment (*p* > 1.05 x 10^−5^) and no SNP showed Linkage Disequilibrium (*k* = 1.12 x 10^7^, *p* < 4.46 x 10^−9^). A total of 61 potential loci under selection was identified and three plastid loci were found, resulting in a dataset with 4,508 SNPs. For effective population size estimates (*N*_e_), a total of 5,100 SNPs were identified. However, 352 SNPs deviated from H-W equilibrium after Bonferroni adjustment (*p* > 9.8 x 10^−6^) and no SNP showed Linkage Disequilibrium (k = 1.3 x 10^7^, *p* < 3.84 x 10^−9^). Seven plastid loci were identified and no loci under selection was found, resulting in a dataset with 4,741 SNPs.

### Paternity assignment

Based on 644 SNPs, sixteen seedlings were assigned to 11 parents within the RNV locality, with some resulting from selfing. However, the parents of five seedlings could not be identified. In RBS, 18 seedlings were assigned to 12 parents within the same locality, but 11 seedlings either had unidentified parents or the inferred probability was less than 95%.

Figure 2 illustrates the dispersal distances of seeds and pollen. In both localities, the majority of dispersal occurred over short distances. For RNV, the average seed dispersal distance was 13.2 meters (m) ± 12.7 SE (range 2.9 - 55 m), while the average pollen dispersal distance was 44.7 m ± 46.4 SE (range 4.25 - 149.1 m). Conversely, in RBS, the average seed dispersal distance was 8.39 m ± 5.7 SE (range 2.3 - 23.6 m), and the average pollen dispersal distance was 33.4 m ± 10.3 SE (range 21.4 - 48 m).

**FIGURE 1.**
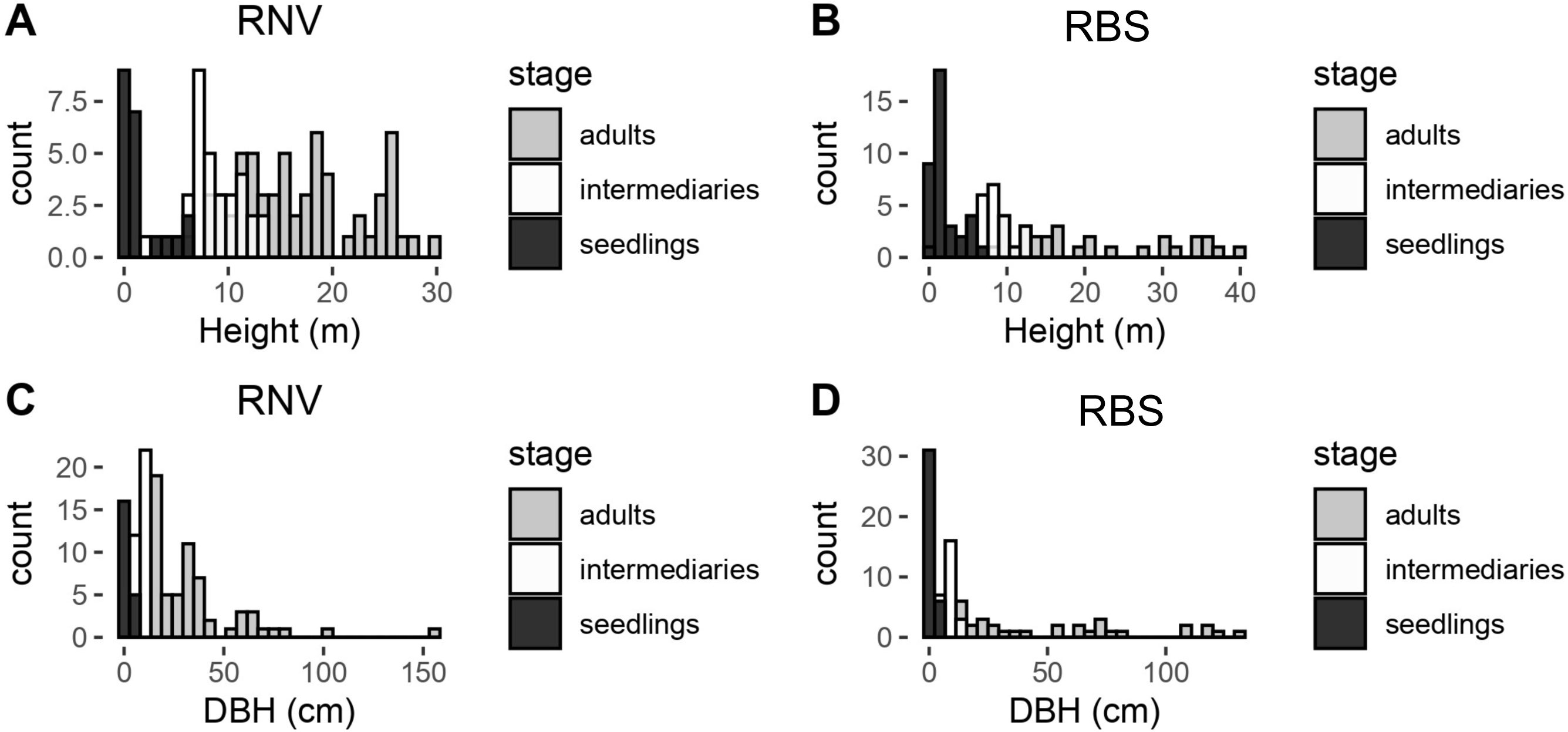
Histograms of the diameter at breast height (DBH) and height distribution for different ontogenetic stages in *Dinizia jueirana-facao* G. P. Lewis & G. S. Siqueira in two Atlantic forest fragments – Reserva Natural da Vale (RNV) and Reserva Biológica de Sooretama (RBS) – located in the Espírito Santo state, southeast Brazil.

**FIGURE 2.**
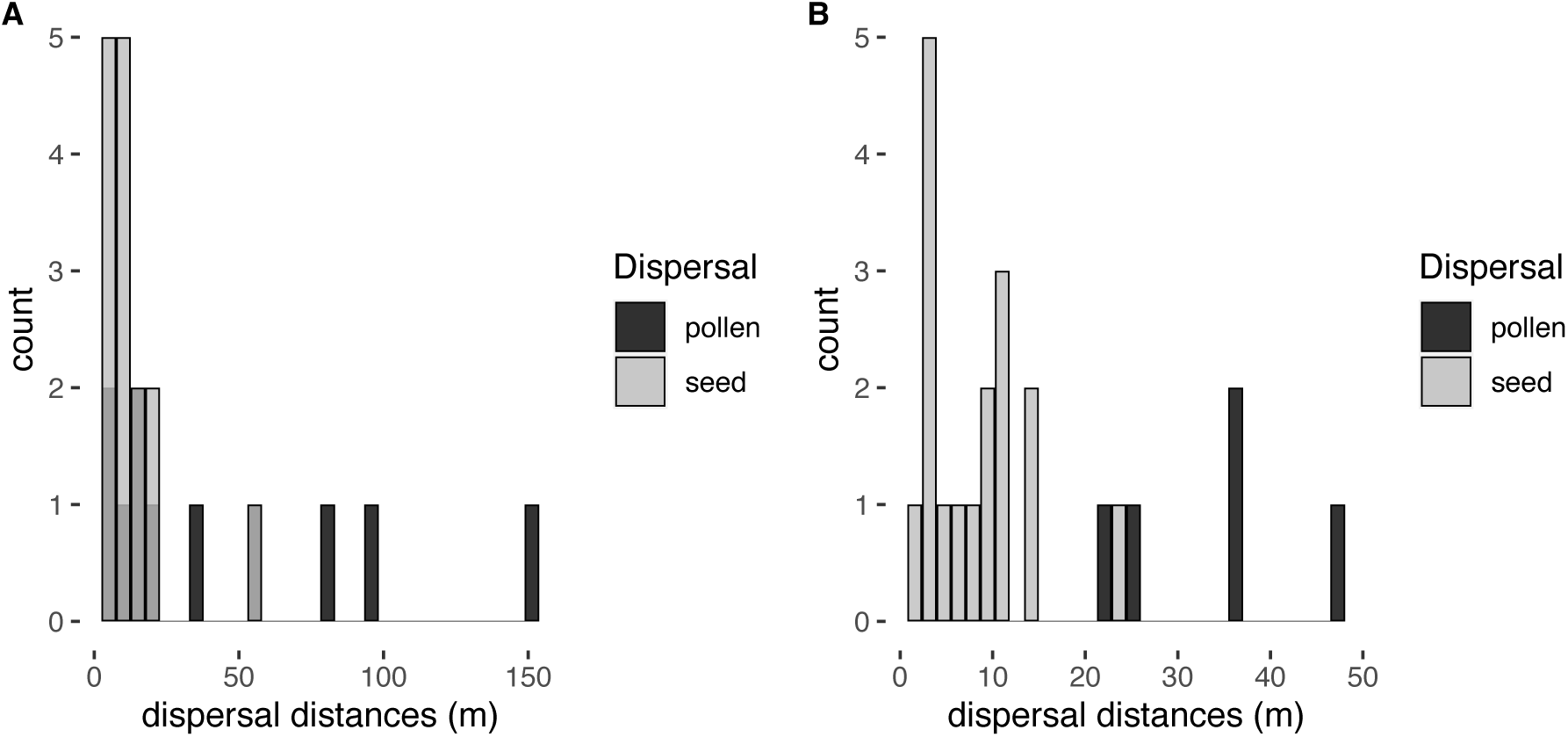
Histogram of gene flow distances for pollen (black bars) and seeds (gray bars) of *Dinizia jueirana-facao* G. P. Lewis & G. S. Siqueira in two Atlantic forest fragments – Reserva Natural da Vale (RNV - A) and Reserva Biológica de Sooretama (RBS - B) – located in the Espírito Santo state, southeast Brazil.

The immigration rates of RNV were determined to be 23.80% for seeds (*m_s_*) and 19.04% for pollen (*m_p_*), whereas in RBS, the rates were 20.68% *m_s_* and 44.82% *m_p_*. Taking into account the variances in pollen and seed dispersal, the total gene flow was estimated to be 34.04 m and 8.38 m for RNV and RBS, respectively.

### Spatial genetic structure and historical dispersal distance

A significant spatial genetic structure, as indicated by the 95% CI, was observed in the RNV locality, with a maximum distance of 134.5 m for seedlings and 53.6 m for reproductive adults. Similarly, significant values were found in RBS, with distances of up to 32.1 m for seedlings and 37.1 m for reproductive adults. Overall, the *F_L_* values decrease and become non-significant in the upper distance classes (Figure 3).

**FIGURE 3.**
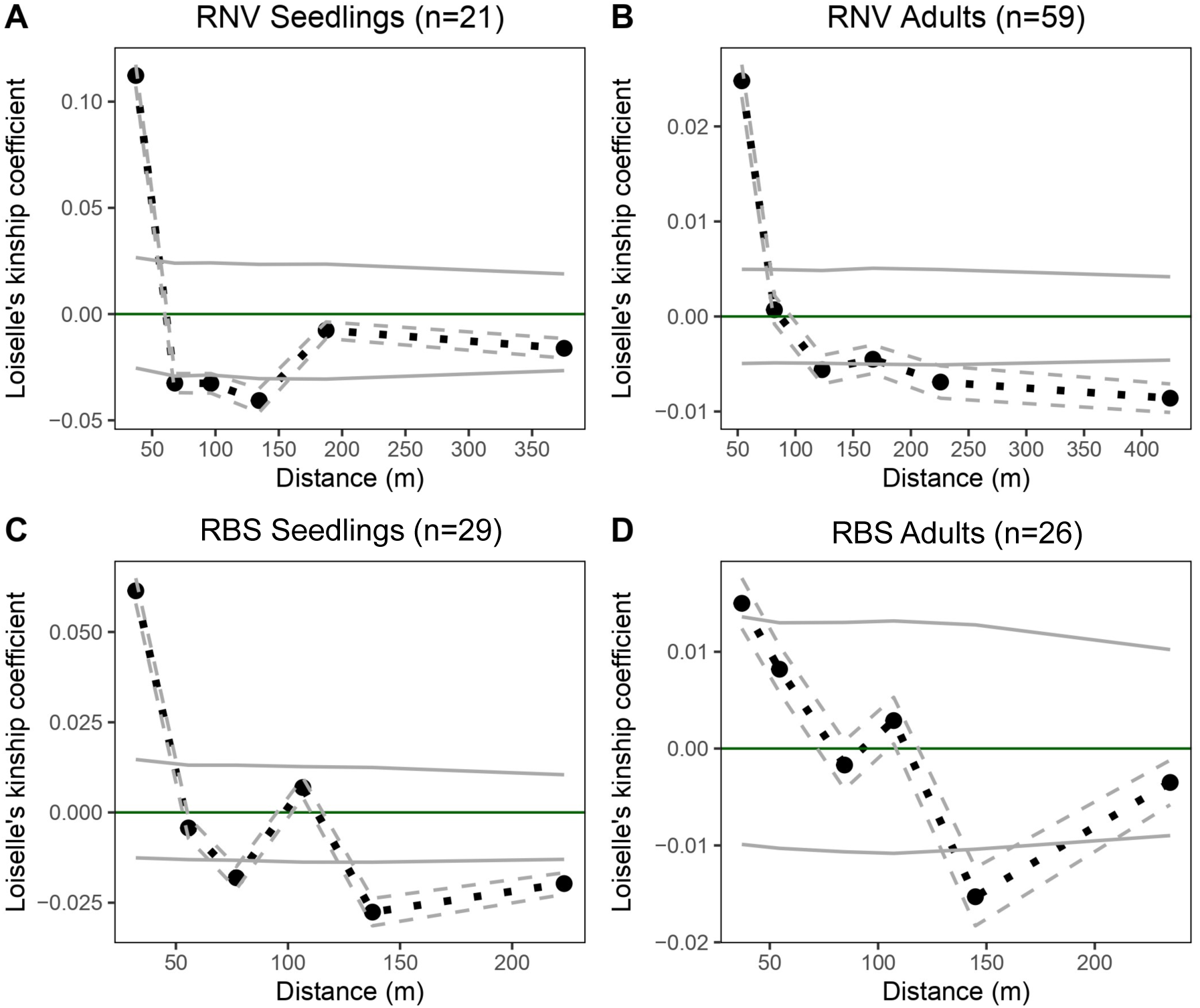
Average Loiselle’s kinship coefficient *F_L_* plotted against geographical distance (black dashed line) for seedlings and adults of *Dinizia jueirana-facao* G. P. Lewis & G. S. Siqueira in two Atlantic forest fragments – Reserva Natural da Vale (RNV) and Reserva Biológica de Sooretama (RBS). The error bars (gray dashed lines) were generated by jackknifing over loci to approximate SE, and 95% CI were generated through 10,000 data permutations and are indicated by gray solid lines.

Both *D. jueirana-facao* reproductive adults and seedlings exhibited variable kinship coefficients across distance classes for both localities, with the highest Loiselle’s kinship coefficient^61^ observed in the first distance class (0-50 m). In RNV, the highest kinship for seedlings (*F_L_*= 0.112, *p* < 0.05) corresponded within theoretical expectations to half-siblings (*F_L_*= 0.125), while in RBS this coefficient (*F_L_*= 0.0614, *p* < 0.05) corresponded to first cousins (*F_L_*= 0.0625). Among *D. jueirana-facao* reproductive adults, the highest kinship (*F_L_*= 0.0248, *p* < 0.05) corresponded within theoretical expectations to full-siblings (*F_L_*= 0.250), while in RBS this coefficient represents half- siblings (*F_L_*= 0.125) for the highest observed estimate (*F_L_*= 0.0150, *p* < 0.05).

The calculated values of *De* for RNV were 0.2 ind.ha^-1^ for adults and 0.48 ind.ha^-1^ for seedlings, while for RBS these values were 0.4 ind.ha^-1^ for adults and 0.44 ind.ha^-1^ for seedlings. The magnitude of SGS, as measured by the *Sp*-statistic, was stronger in RNV locality compared to RBS. For the former locality, the *Sp* values were 0.05088 (CI = 0.05085 - 0.05091) and 0.01664 (CI = 0.01664 - 0.01664) for seedlings and adults, respectively. In RBS, the *Sp* values were 0.03158 (CI = 0.03157 - 0.03159) and 0.01162 (CI = 0.0116 - 0.01162) for seedlings and adults, respectively. Additionally, the SGS was found to be stronger in seedlings compared to adults (Table 1). Considering the variances in pollen and seed dispersal, the total indirect gene flow distance was higher than the contemporary estimates in both localities (Table 1).

**TABLE 1.** Estimates of fine-scale spatial genetic structure (SGS) and of historical dispersal distance for seedlings and reproductive adult trees of *Dinizia jueirana-facao* G. P. Lewis & G. S. Siqueira in two Atlantic forest fragments – Reserva Natural da Vale (RNV) and Reserva Biológica de Sooretama (RBS) – located in the Espírito Santo state, southeast Brazil. The number of individuals and the parameters such as inbreeding coefficient (*F*), average kinship coefficient (*F*_1_) between individuals for the first distance class (i.e., the smallest distance class, which includes distances < 50 m) and its standard error (SE), the *Sp*-statistic, the neighborhood size (*Nb*) and its 95% CI, and the root- mean-squared dispersal distance (σ) and its 95% CI, are shown considering the effective densities of 0.2 for reproductive adult trees and 0.48 for seedlings in RNV, and 0.4 for reproductive adult trees and 0.44 for seedlings in RBS.

| | | $n$ | SGS parameters | | | | Gene dispersal estimates | |
| --- | --- | --- | --- | --- | --- | --- | --- | --- |
| | | | $F$ | $F_1$ | SE | $Sp$ | $Nb$ | $\sigma$ (m) |
| RNV | Seedlings | 21 | 0.108 | 0.112 | 0.005 | 0.051 | 19.65<br>(19.64 - 19.67) | 105.12<br>(105.09 - 105.15) |
|  | Adults | 59 | 0.005 | 0.024 | 0.001 | 0.017 | 60.09<br>(60.08 - 60.1) | 308.09<br>(308.08 - 308.12) |
| RBS | Seedlings | 29 | 0.022 | 0.061 | 0.003 | 0.032 | 31.66<br>(31.65 - 31.68) | 123.4<br>(123.38 - 123.43) |
|  | Adults | 26 | -0.041 | 0.015 | 0.002 | 0.012 | 86.09<br>(86.06 - 86.13) | 192.67<br>(192.63 - 192.7) |

### Mating system, genetic diversity, and inbreeding estimates

Based on the analysis of the mating system, the outcrossing rate was found to be 0.977 (SE±0.004) for RNV and 0.942 (SE± 0.009) for RBS. In terms of genetic variation, the mean allelic richness (*A*) values differed across generations for RNV, with reproductive adults [*A*= 1.711 (95% CI= 1.705 - 1.717)] presenting higher values compared to seedlings [*A*= 1.661 (95% CI= 1.655 - 1.667)]. However, for RBS, there were no significant differences between reproductive adults [*A*= 1.690 (95% CI= 1.684 - 1.696)] and seedlings [*A*= 1.694 (95% CI= 1.689 - 1.700)]. When we compare allelic richness between different localities, we observe that reproductive adults in RNV had higher values than RBS (see Table S7). Nonetheless, seedlings in RNV showed lower values than those observed in RBS.

In the RNV locality, the comparison of average private allelic richness (*R*) between ontogenetic stages showed higher values for reproductive adults [*R*= 0.197 (95% CI= 0.192 – 0.202)] than seedlings [*R*= 0.164 (95% CI= 0.160 – 0.168)]. In contrast, RBS displayed smaller *R* values for reproductive adults [*R*= 0.1766 (95% CI= 0.172 - 0.181)] compared to seedlings [*R*= 0.197 (95% CI= 0.193 - 0.201)]. The results also indicated that reproductive adults in RNV had lower *R* values than those observed in RBS, and the same pattern was observed for seedlings (see Table S7). Lastly, estimates of contemporary *N*_e_ were 40.0 (95% CI = 33.5 - 44.1) for RNV locality and 31.3 (95% CI = 24.6 - 37.7) for RBS.

Regarding the average inbreeding coefficient *F_IS_*, both localities showed negative but not significant values for adults (RNV= -0.051, RBS= -0.049). In addition, seedlings exhibited positive but not significant values (RNV= 0.044, RBS= 0.006). Nevertheless, no statistically significant differences (*p*-value > 0.05) were found when comparing the values between adults and seedlings.

### Population size and demographic structure

A total of 211 individuals were surveyed in the *D*. *jueirana-facao* localities, with 116 individuals in RNV and 95 individuals in RBS. The population density was 2.7 ind.ha^-1^ in RNV and 1.46 ind.ha^-1^ in RBS. Using the previously defined ontogenetic stages, we estimated that in RNV, 18.1% (n=21) of the individuals were seedlings, 29.3% (n=34) were intermediates, and 52.6% (n=61) were classified as adults. In RBS, 38.9% (n=37) were seedlings, 27.4% (n=26) intermediates, and 33.7% (n=32) were identified as adults.

For RNV and RBS, data of DBH and tree height did not exhibit a normal distribution (Table S3). When compared between localities, we found no significant differences in both diameter and height distributions among ontogenetic stages (*p* > 0.05, Table S3), except between RNV adults that showed higher values than RBS adults (*p* < 0.05; Table S3, Figure S3). Graphical analyses of DBH classes revealed a reverse J-shaped size class distribution, accounting for 47.4% (RNV) and 66.3% (RBS) of individuals in the smaller diameter classes (Figure 1). The height distribution showed that 73.2% of individuals in RNV were between 0.08 and 15 m in height, compared to 81% in RBS. In terms of regeneration status, RNV exhibited a fair status, while RBS showed a good status (Table S4).

## DISCUSSION

This study aimed to investigate the genetic and ecological consequences of forest fragmentation on *D. jueirana-facao*, a rare and critically endangered tree species endemic to the Brazilian Atlantic Forest. Although forest fragmentation is commonly associated with reduced genetic diversity, limited gene flow, and altered demographic structures in plant populations^9,11,26,29,73–78^, our findings suggest that *D. jueirana-facao* seems to be resilient to forest fragmentation and habitat loss, as evidenced by a healthy demographic size structure, absence of inbreeding, no apparent reduction in genetic diversity, and high levels of gene flow in the studied areas. We will now discuss the implications of these results, exploring potential mechanisms behind the species’ resilience and factors contributing to genetic diversity and connectivity in the face of forest fragmentation. Additionally, we highlight the implications for conservation and management of *D. jueirana-facao*, emphasizing the need for a comprehensive understanding of its ecological and evolutionary dynamics in the context of ongoing habitat loss and fragmentation.

Our significant discovery of over 211 *D. jueirana-facao* individuals, compared to fewer than 25 reported in a previous study^36^, provides a more accurate assessment of the species’ population abundance. Although the forest fragments where *D. jueirana-facao* occurs are surrounded by roads and an agricultural matrix, the potential presence of individuals outside the study area remains unknown, warranting future field expeditions to enhance our understanding of the species’ geographic distribution and to identify priority sites for conservation efforts.

Contrary to expectations for a threatened plant species^74,75,77,79–81^, the population size structure of *D. jueirana-facao* exhibited a typical reverse J-shaped distribution^82^. This pattern indicates a higher number of individuals in smaller diameter classes^33,83^ and suggests healthy populations that naturally regenerate through recruitment^32,33^. Similar findings have been reported in various plant species, even in disturbed environments^83–85^. However, it is crucial to consider that factors such as resource availability, competition among individuals, natural or anthropogenic disturbances, natural selection against inbreeding, and habitat traits can influence the diameter-height size distribution, as well as seedling recruitment and survival^74,86–88^.

The regeneration status between *D. jueirana-facao* localities showed different patterns, with RNV showing fair regeneration, while RBS exhibited good regeneration. This is a positive aspect for the species, as the presence of seedlings in populations may be attributed to unlimited abiotic factors and a positive trade-off between predation, selection and/or competition and survival rate^33,85^. Overall, our results demonstrate a healthy demographic size structure for *D. jueirana-facao*, indicating its resilience to environmental disturbances. However, long-term monitoring of *D. jueirana-facao* localities, collecting annual demographic data, is imperative not only for the protection of their populations but also to establish a comprehensive understanding of population dynamics at an ecological level.

Contemporary estimates of gene flow in both investigated localities were lower than historical estimates, highlighting the threat posed by forest fragmentation in the species *D. jueirana-facao*. This pattern was particularly evident in RBS, where even lower values were observed. These findings are consistent with previous studies involving trees^22,31,89,90^. However, despite the occurrence of disturbances, gene dispersal can persist at significant levels within populations^30,91^, indicating the existence of mechanisms promoting genetic connectivity at restricted geographic scales. Therefore, the variations of contemporary gene flow can be attributed to a range of factors, such as population density, mating system, and specific responses to anthropogenic disturbances due to the range of distinct plant life-history traits^92–95^.

Regarding the gene dispersal distances of *D. jueirana-facao*, contemporary estimates reveal that gene flow by pollen is remarkably more efficient, with distances approximately four times greater than seed dispersal in the RBS locality, as previously observed for RNV^31^. This trend is consistent with previous studies in tree species^55,94,96,97^. This highlights the critical role of pollen flow to prevent inbreeding and self-pollination in tree species with both hermaphrodite and functional male flowers, such as *D. jueirana-facao*. In fact, the pattern of pollen flow observed for *D. jueirana-facao* seems to be directly linked to the lack of inbreeding (i.e., inbreeding rates were close to zero) and very low historical selfing rates.

The effectiveness of pollen or seed dispersal contributes to species resilience against the negative effects of forest fragmentation^98^. For instance, in fragmented landscapes, gene flow through pollen can increase, as observed in tropical species with specialized pollination systems^99,100^ or even for plant species with generalized pollination systems such *as D. excelsa*, where fragmentation led to changes in pollinators allowing pollen transport over very long distances^39^. Other studies have shown the positive effect of secondary vegetation in reducing the average isolation of forest fragments, enabling greater landscape connectivity, improving the mobility of potential dispersers, and maintaining gene flow^101–103^. Nevertheless, some studies have reported that even wind-pollinated tree species can be negatively affected by fragmentation and habitat loss^104–107^, indicating that knowledge of the species’ life-history traits is the main, but not the only factor in predicting the genetic consequences of habitat fragmentation.

The primary means of seed dispersal for *D. jueirana-facao* is gravity, while pollen may potentially be dispersed by wind. This mode of pollen dispersal may contribute to its efficient long-distance distribution. In addition, the open landscape may be contributing to long-distance pollen flow, which could mitigate the potential negative genetic effects of forest fragmentation. Indeed, we observed gene immigration beyond the forest fragment edges, indicating long-distance pollen movement. However, it is important to consider the estimated immigration rates with caution due to uncontrolled factors in the sampling, such as missing genotypes caused by tree mortality and illegal timber activities in the past years^91,108^.

Despite the presence of gene immigration, we did not identify any parentage assignments between the two studied localities, as the assignments were exclusive to each locality. While the population census was conducted in RNV locality, we covered approximately 80% of the locality area. This suggests that the parent candidates in the RBS locality may not have been sampled or may have been missing due to the presence of dead and decaying trees. Nevertheless, our results indicate that the RNV locality is not genetically isolated, as there is evidence of long-distance pollen dispersal between the forest fragment and the nearest pollen source, as previously observed by Nazareno and Knowles^31^.

Considering the uncertainty surrounding the genetic connectivity of RBS locality, and even though forest fragmentation has no apparent negative effects on genetic diversity, it is crucial to promote connectivity among the remaining fragments. This will help avoid the negative genetic effects observed in small populations^11,109–111^. Strategies such as implementing ecological corridors could effectively minimize further detrimental genetic and ecological effects in these small localities^112,113^.

In terms of historical gene flow estimates, we observed SGS in both reproductive adults and seedlings of both localities. This indicates a high probability of relatedness between individuals (e.g., up to 134.5 m in RNV seedlings), with a decline as geographic distances increase. Typically, plant species with long-distance dispersal of pollen and seeds exhibit low or no SGS^114^. Our study reveals a significant SGS, with higher *Sp* values (*Sp* range = 0.01162 - 0.05088) compared to other tree species [e.g., *Parkia biglobosa* (*Sp* range = 0.0002 to 0.003^90^), *Entandrophragma cylindricum* (*Sp* = 0.0058^89^)]. This suggests that the observed SGS in *D. jueirana-facao* may be a result of seed dispersal near mother trees, with pollen dispersal playing a secondary role. Previous studies have shown that the strength of SGS is influenced by pollination and seed dispersal vectors^60,95,114–117^. For instance, Vekemans and Hardy^60^ and Dick et al.^115^ demonstrated that plants dispersed by gravity tend to have higher *Sp* values compared to species dispersed by animals, even if pollen movement is random and long-distance^115^.

Despite the expected loss of genetic variation and reduced gene flow in fragmented and small plant populations^9,11,26,29,73,76^, our results seemingly support the hypothesis that *D. jueirana-facao* does not display measurable effects resulting from forest fragmentation. Consistent with the findings for *D. jueirana-facao*, numerous studies, including meta-analyses, have reported a perceived resilience of tree species to the population genetic effects of habitat loss and fragmentation^20,90,118–122^. This unexpected scenario arises from the long lifespan of trees, the presence of overlapping generations that help prevent the loss of genetic diversity, the flexible mating systems in some species that can bypass self-incompatibility and produce self-pollinated offspring, and the prevalence of extensive gene flow. Indeed, the extensive pollen flow over long distances and migration events, coupled with our contrasting findings of genetic diversity between the two *D. jueirana-facao* localities, indicate that forest fragmentation has not yet hurt this tree species.

Although we did not find evidence of inbreeding in either locality or for the two ontogenetic stages of *D. jueirana-facao*, our results reveal a slight reduction in allelic and private allelic richness only for RNV. Undoubtedly, allelic richness is the genetic parameter most sensitive to the effects of forest fragmentation compared to other commonly used metrics, making it suitable for tracking recent effects of habitat fragmentation^104,123,124^. While reduced genetic diversity associated with forest fragmentation has been well-documented in many other species^19,87,125–127^, it is also common for genetic diversity to be maintained^20,119,120,128^. This contrast reveals how life-history traits (e.g., life cycle, growth form, mating system) strongly influence the amount and distribution of genetic variation in natural populations^121,129–131^.

While historical outcrossing rates suggest an allogamous mating system for *D. jueirana-facao*, paternity assignments have revealed a higher incidence of contemporary selfing events in the RNV locality compared to RBS (see Table S8). Indeed, it was in the RNV locality that we found a slight loss on genetic variation. It is worth noting that outcrossing rates can vary spatially and temporally, and the prevalence of different mating systems can be influenced by abiotic factors that affect plant fitness^132,133^. For instance, despite expectations of autogamous and apomictic systems in anthropized environments, Silva et al.^134^ found a higher occurrence of allogamous species. Landscape anthropization^135^, environmental changes, plant density, and demographic size structure can all impact mating rates^132^. Therefore, conducting long-term studies related to genetic, phenology and pollination biology are crucial to better understand the factors that affect reproduction in *D. jueirana-facao*.

This study offers valuable insights for the conservation of *D. jueirana-facao*. Specifically, our estimates of indirect dispersal indicate the need to collect seeds from reproductive trees separated by at least 134.5 m for RNV and 37.1 m for RBS. This information should be considered in ex-situ conservation and restoration programs^31,136–139^. It is important to note that, while the individuals in RNV and RBS are located in protected areas, with an apparent lacking of inbreeding and showing a healthy demographic size structure, the RBS locality is threatened by ongoing habitat loss due to deforestation associated with human activities such as agriculture, livestock farming, and mining. Therefore, continuous monitoring of the localities where *D. jueirana-facao* occurs is urgently needed as part of conservation programs. Particularly, agriculture in and around these forest fragments and ecotourism activities should be addressed in collaboration with stakeholders (e.g., local communities and important national companies such as Vale do Rio Doce) to ensure the maintenance and conservation of this rare genetic resource. Additionally, based on estimates of contemporary *N*_e_, the findings indicate that *D. jueirana-facao* have *N*_e_ values below the recommended threshold of 50 set by the "50/500 rule" for short-term inbreeding concerns. The species also does not meet the minimum *N*_e_ threshold of 500^29,140^, or as proposed by Frankham^141^, a *N*_e_ value greater than 1000, for maintaining long-term evolutionary potential. This suggests that both localities may be at risk of reduced genetic diversity and increased susceptibility to genetic problems in the future. Therefore, keeping the areas where *D. jueirana-facao* occur without further degradation and habitat loss would help to maintain more than 97% of *D. jueirana-facao*’s genetic diversity – an amount above that recommended for the CBD draft post-2020 global biodiversity framework^124,141^.

While our study aimed to assess the genetic and ecological effects of forest fragmentation in a rare, endemic, and critically endangered tree species, there are limitations and caveats to consider regarding the contemporary gene flow analyses, particularly related to sampling design (e.g., incomplete sampling in the RBS locality) and limited knowledge about the presence of *D. jueirana-facao* outside the two known localities. Although the isolation of the RBS locality remains uncertain due to unsampled or missing candidate parents (e.g., decayed or removed trees due to illegal timber activity), our study provides evidence that this locality does not exhibit the expected characteristics of isolated and small populations affected by forest fragmentation^19,22,127^. The fragmented populations of *D. jueirana-facao* in RBS demonstrates remarkable resilience to cope with the effects of forest fragmentation. Although our results are promising, long-term ecological and genetic studies are still needed. In addition to our SNP-based approach, deeper genomic analyses such as the identification and characterization of structural variants – which has been proposed for threatened species^142^– would provide an excellent opportunity to enhance the conservation of *D. jueirana-facao*. These complementary approaches should be prioritized as a foundation for conservation and management programs for *D. jueirana-facao*.

## ACKNOWLEDGEMENTS

We thank Domingos A. Folli for assistance during fieldwork. We also thank Reserva Natural Vale for field support.

## DATA ACCESSIBILITY

SNP datasets for *D. jueirana-facao* are available for download from the FigshareDigital Repository (https://figshare.com/s/6df6cc1f3c5b47306804, https://doi.org/10.6084/m9.figshare.24925635). The raw data generated for *D. jueirana-facao* are available to download from the ENA (European Nucleotide Archive) under accession number ERP129560.

## AUTHOR CONTRIBUTIONS

AGN designed and obtained project funds, collected samples, and conducted molecular lab work. TMT collected samples and provided analytical support. KASP performed analyses, and led the writing of the manuscript with input from AGN and TMT. All authors approved the submitted version of this manuscript.

## REFERENCES

1. Taubert, F. et al. Global patterns of tropical forest fragmentation. Nature 554, 519–522 (2018).

2. Fischer, R. et al. Accelerated forest fragmentation leads to critical increase in tropical forest edge area. Sci. Adv. 7, eabg7012 (2021).

3. Ribeiro, M. C., Metzger, J. P., Martensen, A. C., Ponzoni, F. J. & Hirota, M. M. The Brazilian Atlantic Forest: How much is left, and how is the remaining forest distributed? Implications for conservation. Biol. Conserv. 142, 1141–1153 (2009).

4. Rezende, C. L. et al. From hotspot to hopespot: An opportunity for the Brazilian Atlantic Forest. *Perspect*. Ecol. Conserv. 16, 208–214 (2018).

5. Fundação SOS Mata Atlântica & INPE. Atlas dos Remanescentes Florestais da Mata Atlântica. Período 2021 - 2022. (2023).

6. da Fonseca, G. A. B. The vanishing Brazilian Atlantic forest. Biol. Conserv. 34, 17–34 (1985).

7. Diniz, M. F., Coelho, M. T. P., Sánchez-Cuervo, A. M. & Loyola, R. How 30 years of land-use changes have affected habitat suitability and connectivity for Atlantic Forest species. Biol. Conserv. 274, 109737 (2022).

8. Saunders, D. A., Hobbs, R. J. & Margules, C. R. Biological Consequences of Ecosystem Fragmentation: A Review. Conserv. Biol. 5, 18–32 (1991).

9. Young, A., Boyle, T. & Brown, T. The population genetic consequences of habitat fragmentation for plants. Trends Ecol. Evol. 11, 413–418 (1996).

10. Honnay, O. & Jacquemyn, H. Susceptibility of Common and Rare Plant Species to the Genetic Consequences of Habitat Fragmentation. Conserv. Biol. 21, 823–831 (2007).

11. Schlaepfer, D. R., Braschler, B., Rusterholz, H. & Baur, B. Genetic effects of anthropogenic habitat fragmentation on remnant animal and plant populations: a meta-analysis. Ecosphere 9, e02488 (2018).

12. Horváth, Z., Ptacnik, R., Vad, C. F. & Chase, J. M. Habitat loss over six decades accelerates regional and local biodiversity loss via changing landscape connectance. Ecol. Lett. 22, 1019–1027 (2019).

13. Leigh, D. M., Hendry, A. P., Vázquez-Domínguez, E. & Friesen, V. L. Estimated six per cent loss of genetic variation in wild populations since the industrial revolution. Evol. Appl. 12, 1505–1512 (2019).

14. Hansen, M. C. et al. The fate of tropical forest fragments. Sci. Adv. 6, eaax8574 (2020).

15. Chiriboga-Arroyo, F. et al. Genetic threats to the Forest Giants of the Amazon: Habitat degradation effects on the socio-economically important Brazil nut tree (*Bertholletia excelsa*). Plants People Planet 3, 194–210 (2021).

16. Cristóbal-Pérez, E. J., Fuchs, E. J., Martén-Rodríguez, S. & Quesada, M. Habitat fragmentation negatively affects effective gene flow via pollen, and male and female fitness in the dioecious tree, *Spondias purpurea* (Anacardiaceae). Biol. Conserv. 256, 109007 (2021).

17. Łabiszak, B., Zaborowska, J., Wójkiewicz, B. & Wachowiak, W. Molecular and paleo-climatic data uncover the impact of an ancient bottleneck on the demographic history and contemporary genetic structure of endangered *Pinus uliginosa*. J. Syst. Evol. 59, 596–610 (2021).

18. Obico, J. J. A. et al. No Evidence of Low Genetic Diversity Despite High Levels of Inbreeding and Poor Genetic Connectivity Among *Tetrastigma loheri* (Vitaceae) Populations in Remaining Forest Areas in Cebu, Philippines. Syst. Bot. 46, 951–961 (2021).

19. Waqar, Z. et al. Gene Flow and Genetic Structure Reveal Reduced Diversity between Generations of a Tropical Tree, *Manilkara multifida* Penn., in Atlantic Forest Fragments. Genes 12, 2025 (2021).

20. De Santana, N. S. et al. Genetic resilience of Atlantic forest trees to impacts of biome loss and fragmentation. Eur. J. For. Res. 142, 161–174 (2023).

21. Exposito-Alonso, M. et al. Genetic diversity loss in the Anthropocene. Science 377, 1431–1435 (2022).

22. Jiang, Q., Xu, Q., Pan, J., Yao, X. & Cheng, Z. Impacts of Chronic Habitat Fragmentation on Genetic Diversity of Natural Populations of ’*Prunus persica* in China. Plants 11, 1458 (2022).

23. Zhang, L. et al. Assessing genetic diversity in critically endangered *Chieniodendron hainanense* populations within fragmented habitats in Hainan. Sci. Rep. 14, 6988 (2024).

24. Fisher, R. The Genetical Theory of Natural Selection. (1930).

25. Hughes, A. R., Inouye, B. D., Johnson, M. T. J., Underwood, N. & Vellend, M. Ecological consequences of genetic diversity. Ecol. Lett. 11, 609–623 (2008).

26. Aguilar, R., Quesada, M., Ashworth, L., Herrerias-Diego, Y. & Lobo, J. Genetic consequences of habitat fragmentation in plant populations: susceptible signals in plant traits and methodological approaches. Mol. Ecol. 17, 5177–5188 (2008).

27. Vranckx, G., Jacquemyn, H., Muys, B. & Honnay, O. Meta-Analysis of Susceptibility of Woody Plants to Loss of Genetic Diversity through Habitat Fragmentation. Conserv. Biol. 26, 228–237 (2012).

28. Tambarussi, E. V., Boshier, D., Vencovsky, R., Freitas, M. L. M. & Sebbenn, A. M. Inbreeding depression from selfing and mating between relatives in the Neotropical tree *Cariniana legalis* Mart. Kuntze. Conserv. Genet. 18, 225–234 (2017).

29. Allendorf, F. W. et al. Conservation and the Genomics of Populations. (Oxford University Press, 2022).

30. Oddou-Muratorio, S. & Klein, E. K. Comparing direct vs. indirect estimates of gene flow within a population of a scattered tree species. Mol. Ecol. 17, 2743–2754 (2008).

31. Nazareno, A. G. & Knowles, L. L. There Is No ‘Rule of Thumb’: Genomic Filter Settings for a Small Plant Population to Obtain Unbiased Gene Flow Estimates. Front. Plant Sci. 12, 677009 (2021).

32. Boz, G. & Maryo, M. Woody Species Diversity and Vegetation Structure of Wurg Forest, Southwest Ethiopia. Int. J. For. Res. 2020, 8823990 (2020).

33. Balemlay, S. & Siraj, M. Population Structure and Regeneration Status of Woody Species in Kenech Forest, Southwest Ethiopia. Int. J. For. Res. 2021, 6640285 (2021).

34. Kierulff, M. C. M., Avelar, L. D. S., Ferreira, M. dos S., Povoa, K. F. & Bérnils, R. S. Reserva Natural Vale: história e aspectos físicos. Ciênc. Ambiente 49, 7–40 (2014).

35. Rolim, S., et al. Floresta Atlântica de tabuleiro: diversidade e endemismos na Reserva Natural da Vale. Rupestre (2016).

36. Lewis, G. P., Siqueira, G. S., Banks, H. & Bruneau, A. The majestic canopy-emergent genus *Dinizia* (Leguminosae: Caesalpinioideae), including a new species endemic to the Brazilian state of Espírito Santo. Kew Bull. 72, 48 (2017).

37. IUCN. (2017). The IUCN Red List of Threatened Species. Version 2022–2. https://www.iucnredlist.org. Accessed on July 18, 2023.

38. Brasil. Portaria MMA N° 148, de 7 de junho de 2022. Dispõem sobre a atualização da Lista Nacional de Espécies Ameaçadas de Extinção. Diário Oficial da União. 74 (2022).

39. Dick, C. W., Etchelecu, G. & Austerlitz, F. Pollen dispersal of tropical trees (*Dinizia excelsa* : Fabaceae) by native insects and African honeybees in pristine and fragmented Amazonian rainforest. Mol. Ecol. 12, 753–764 (2003).

40. Milios, E., Kitikidou, K. G., Dalakouras, V. & Pipinis, E. Diameter At Breast Height Estimated From Stumps In *Quercus frainetto* In The Region Of Evros In Northeastern Greece. CERNE 22, 337–344 (2016).

41. Peterson, B. K., Weber, J. N., Kay, E. H., Fisher, H. S. & Hoekstra, H. E. Double Digest RADseq: An Inexpensive Method for De Novo SNP Discovery and Genotyping in Model and Non-Model Species. PLoS ONE 7, e37135 (2012).

42. Nazareno, A. G., Bemmels, J. B., Dick, C. W. & Lohmann, L. G. Minimum sample sizes for population genomics: an empirical study from an Amazonian plant species. Mol. Ecol. Resour. 17, 1136–1147 (2017).

43. Andrews, S. FastQC A Quality Control tool for High Throughput Sequence Data. Babraham Bioinformatics (2010).

44. Catchen, J. M., Amores, A., Hohenlohe, P., Cresko, W. & Postlethwait, J. H. *Stacks* : Building and Genotyping Loci *De Novo* From Short-Read Sequences. G3 GenesGenomesGenetics 1, 171–182 (2011).

45. Catchen, J., Hohenlohe, P. A., Bassham, S., Amores, A. & Cresko, W. A. Stacks: an analysis tool set for population genomics. Mol. Ecol. 22, 3124–3140 (2013).

46. Rochette, N. C., Rivera-Colón, A. G. & Catchen, J. M. Stacks 2: Analytical methods for paired-end sequencing improve RADseq-based population genomics. Mol. Ecol. 28, 4737–4754 (2019).

47. Hohenlohe, P. A. et al. Population Genomics of Parallel Adaptation in Threespine Stickleback using Sequenced RAD Tags. PLoS Genet. 6, e1000862 (2010).

48. Paradis, E. pegas: an R package for population genetics with an integrated–modular approach. Bioinformatics 26, 419–420 (2010).

49. Rousset, F. GENEPOP ’007: a complete re-implementation of the GENEPOP software for Windows and Linux. Mol. Ecol. Resour. 8, 103–106 (2008).

50. Rice, W. R. Analyzing Tables of Statistical Tests. Evolution 43, 223 (1989).

51. Altschul, S. F., Gish, W., Miller, W., Myers, E. W. & Lipman, D. J. Basic local alignment search tool. J. Mol. Biol. 215, 403–410 (1990).

52. Foll, M. & Gaggiotti, O. A Genome-Scan Method to Identify Selected Loci Appropriate for Both Dominant and Codominant Markers: A Bayesian Perspective. Genetics 180, 977–993 (2008).

53. Wang, J. Sibship Reconstruction From Genetic Data With Typing Errors. Genetics 166, 1963–1979 (2004).

54. Dow, B. D. & Ashley, M. V. Microsatellite analysis of seed dispersal and parentage of saplings in bur oak, *Quercus macrocarpa*. Mol. Ecol. 5, 615–627 (1996).

55. Guidugli, M. C. et al. Small but not isolated: a population genetic survey of the tropical tree *Cariniana estrellensis* (Lecythidaceae) in a highly fragmented habitat. Heredity 116, 339–347 (2016).

56. Crawford, T. J. What is a population? Evol. Ecol. (1984).

57. Smouse, P. E. & Sork, V. L. Measuring pollen flow in forest trees: an exposition of alternative approaches. For. Ecol. Manag. 197, 21–38 (2004).

58. Burczyk, J., Adams, W. T. & Shimizu, J. Y. Mating patterns and pollen dispersal in a natural knobcone pine (*Pinus attenuata* Lemmon.) stand. Heredity 77, 251–260 (1996).

59. Sebbenn, A. M. et al. Low levels of realized seed and pollen gene flow and strong spatial genetic structure in a small, isolated and fragmented population of the tropical tree *Copaifera langsdorffii* Desf. Heredity 106, 134–145 (2011).

60. Vekemans, X. & Hardy, O. J. New insights from fine-scale spatial genetic structure analyses in plant populations. Mol. Ecol. 13, 921–935 (2004).

61. Loiselle, B. A., Sork, V. L., Nason, J. & Graham, C. Spatial genetic structure of a tropical understory shrub, *Psychotria officinalis* (Rubiaceae). Am. J. Bot. 82, 1420–1425 (1995).

62. Peakall, R., Ruibal, M. & Lindenmayer, D. B. Spatial autocorrelation analysis offers new insights into gene flow in the Australian bush rat, *Rattus fuscipes*. Evolution 57, 1182–1195 (2003).

63. Shchipanov, N. A., Artamonov, A. V., Titov, S. V. & Pavlova, S. V. Fluctuating fine-scale spatial genetic structure in the common shrews (Eulipotyphla, Mammalia). Integr. Zool. 18, 469–492 (2023).

64. Hardy, O. J. & Vekemans, X. SPAGeDi: a versatile computer program to analyse spatial genetic structure at the individual or population levels. Mol. Ecol. Notes 2, 618–620 (2002).

65. Szpiech, Z. A., Jakobsson, M. & Rosenberg, N. A. ADZE: a rarefaction approach for counting alleles private to combinations of populations. Bioinformatics 24, 2498–2504 (2008).

66. Do, C. et al. NEESTIMATOR v2: re-implementation of software for the estimation of contemporary effective population size (*N _e_*) from genetic data. Mol. Ecol. Resour. 14, 209–214 (2014).

67. Schwartz, G. et al. Profitability of silvicultural treatments in logging gaps in the Brazilian Amazon. J. Trop. For. Sci. 68–78 (2016).

68. Locosselli, G. M., Krottenthaler, S., Pitsch, P., Anhuf, D. & Ceccantini, G. Age and Growth Rate of Congeneric Tree Species (hymenaea Spp.-Leguminosae) Inhabiting Different Tropical Biomes. Erdkunde 71, 45–57 (2017).

69. Wright, S. ISOLATION BY DISTANCE. Genetics 28, 114–138 (1943).

70. Keenan, K., McGinnity, P., Cross, T. F., Crozier, W. W. & Prodöhl, P. A. diveRsity: An R package for the estimation and exploration of population genetics parameters and their associated errors. Methods Ecol. Evol. 4, 782–788 (2013).

71. R Core Team. R: A Language and Environment for Statistical Computing. R Foundation for Statistical Computing, Vienna. https://www.R-project.org (2018).

72. Baidya, S., Thakur, B. & Devi, A. The influence of tree population structure on regeneration potential in the sacred forests of Assam, India. Trop. Ecol. 63, 239–251 (2022).

73. Lowe, A. J., Boshier, D., Ward, M., Bacles, C. F. E. & Navarro, C. Genetic resource impacts of habitat loss and degradation; reconciling empirical evidence and predicted theory for neotropical trees. Heredity 95, 255–273 (2005).

74. Nazareno, A. G. & Dos Reis, M. S. At Risk of Population Decline? An Ecological and Genetic Approach to the Threatened Palm Species *Butia eriospatha* (Arecaceae) of Southern Brazil. J. Hered. 105, 120–129 (2014).

75. Rother, D. C., Rodrigues, R. R. & Pizo, M. A. Bamboo thickets alter the demographic structure of *Euterpe edulis* population: A keystone, threatened palm species of the Atlantic forest. Acta Oecologica 70, 96–102 (2016).

76. Frankham, R. et al. Genetic Management of Fragmented Animal and Plant Populations. (Oxford University Press Oxford, 2017).

77. Khosa, A. et al. The impact of elephants (*Loxodonta africana*) on the Baobab (*Adansonia digitata*) in a semi-arid savanna. Glob. Ecol. Conserv. 46, e02556 (2023).

78. Wu, Q. et al. Genetic diversity, population genetic structure and gene flow in the rare and endangered wild plant *Cypripedium macranthos* revealed by genotyping-by-sequencing. BMC Plant Biol. 23, 254 (2023).

79. Carvalho, F. A. & Nascimento, M. T. Estrutura diamétrica da comunidade e das principais populações arbóreas de um remanescente de Floresta Atlântica Submontana (Silva Jardim-RJ, Brasil). Rev. Árvore 33, 327–337 (2009).

80. Helm, C. V. & Witkowski, E. T. F. Characterising wide spatial variation in population size structure of a keystone African savanna tree. For. Ecol. Manag. 263, 175–188 (2012).

81. Cousins, S. R., Witkowski, E. T. F. & Pfab, M. F. Elucidating patterns in the population size structure and density of *Aloe plicatilis*, a tree Aloe endemic to the Cape fynbos, South Africa. South Afr. J. Bot. 90, 20–36 (2014).

82. Meyer, H. A. Structure, Growth, and Drain in Balanced Uneven-Aged Forests. J. For. 50, 85–92 (1952).

83. Atsbha, T., Desta, A. B. & Zewdu, T. Woody species diversity, population structure, and regeneration status in the Gra-Kahsu natural vegetation, southern Tigray of Ethiopia. Heliyon 5, e01120 (2019).

84. Muluneh, M. G., Feyissa, M. T. & Wolde, T. M. Effect of forest fragmentation and disturbance on diversity and structure of woody species in dry Afromontane forests of northern Ethiopia. Biodivers. Conserv. 30, 1753–1779 (2021).

85. Yemata, G. & Haregewoien, G. Floristic composition, structure and regeneration status of woody plant species in Northwest Ethiopia. Trees For. People 9, 100291 (2022).

86. Takahashi, K., Ikeyama, Y. & Okuhara, I. Stand dynamics and competition in a mixed forest at the northern distribution limit of evergreen hardwood species. Ecol. Evol. 8, 11199–11212 (2018).

87. Moraes, M. A. et al. Long-distance pollen and seed dispersal and inbreeding depression in *Hymenaea stigonocarpa* (Fabaceae: Caesalpinioideae) in the Brazilian savannah. Ecol. Evol. 8, 7800–7816 (2018).

88. West, P. W. Quantifying effects on tree growth rates of symmetric and asymmetric inter-tree competition in even-aged, monoculture *Eucalyptus pilularis* forests. Trees 37, 239–254 (2023).

89. Monthe, F. K., Hardy, O. J., Doucet, J., Loo, J. & Duminil, J. Extensive seed and pollen dispersal and assortative mating in the rain forest tree *Entandrophragma cylindricum* (Meliaceae) inferred from indirect and direct analyses. Mol. Ecol. 26, 5279–5291 (2017).

90. Lompo, D., Vinceti, B., Konrad, H., Duminil, J. & Geburek, T. Fine-scale spatial genetic structure, mating, and gene dispersal patterns in *Parkia biglobosa* populations with different levels of habitat fragmentation. Am. J. Bot. 107, 1041–1053 (2020).

91. Sola, G., El Mujtar, V., Gallo, L., Vendramin, G. G. & Marchelli, P. Staying close: short local dispersal distances on a managed forest of two Patagonian *Nothofagus* species. For. Int. J. For. Res. 93, 652–661 (2020).

92. Bacles, C. F. E. & Jump, A. S. Taking a tree’s perspective on forest fragmentation genetics. Trends Plant Sci. 16, 13–18 (2011).

93. Hamrick, J. L. & Trapnell, D. W. Using population genetic analyses to understand seed dispersal patterns. Acta Oecologica 37, 641–649 (2011).

94. Hardy, O. J. et al. Seed and pollen dispersal distances in two African legume timber trees and their reproductive potential under selective logging. Mol. Ecol. 28, 3119–3134 (2019).

95. Goncalves, A. L., García, M. V., Barrandeguy, M. E., González-Martínez, S. C. & Heuertz, M. Spatial genetic structure and mating system in forest tree populations from seasonally dry tropical forests: a review. Tree Genet. Genomes 18, 18 (2022).

96. Angbonda, D.-. M. A., Monthe, F. K., Bourland, N., Boyemba, F. & Hardy, O. J. Seed and pollen dispersal and fine-scale spatial genetic structure of a threatened tree species: *Pericopsis elata* (HARMS) Meeuwen (Fabaceae). Tree Genet. Genomes 17, 27 (2021).

97. Sujii, P. S. et al. High gene flow through pollen partially compensates spatial limited gene flow by seeds for a Neotropical tree in forest conservation and restoration areas. Conserv. Genet. 22, 383–396 (2021).

98. Sen, S. & Ravikanth, G. Genetic Consequences of Fragmentation in Tropical Forests: Novel Approaches to Assess and Monitor Critically Endangered Species. in Molecular Genetics and Genomics Tools in Biodiversity Conservation (eds Kumar, A., Choudhury, B., Dayanandan, S. & Khan, M. L.) 79–95 (Springer Nature Singapore, Singapore, 2022).

99. Nason, J. D. & Hamrick, J. L. Reproductive and Genetic Consequences of Forest Fragmentation: Two Case Studies of Neotropical Canopy Trees. J. Hered. 88, 264–276 (1997).

100. Nazareno, A. G. & De Carvalho, D. What the reasons for no inbreeding and high genetic diversity of the neotropical fig tree *Ficus arpazusa*? Conserv. Genet. 10, 1789–1793 (2009).

101. Molin, P. G., Chazdon, R., Frosini De Barros Ferraz, S. & Brancalion, P. H. S. A landscape approach for cost-effective large-scale forest restoration. J. Appl. Ecol. 55, 2767–2778 (2018).

102. Ribeiro, S. E. et al. Do anthropogenic matrix and life-history traits structure small mammal populations? A meta-analytical approach. Conserv. Genet. 22, 703–716 (2021).

103. Alencar, L., Escada, M. I. S. & Camargo, J. L. C. Forest regeneration pathways in contrasting deforestation patterns of Amazonia. *Front*. Environ. Sci. 11, (2023).

104. Jump, A. S. & Peñuelas, J. Genetic effects of chronic habitat fragmentation in a wind-pollinated tree. Proc. Natl. Acad. Sci. 103, 8096–8100 (2006).

105. Barbeta, A., Peñuelas, J., Ogaya, R. & Jump, A. S. Reduced tree health and seedling production in fragmented *Fagus sylvatica* forest patches in the Montseny Mountains (NE Spain). For. Ecol. Manag. 261, 2029–2037 (2011).

106. Kitamura, K. & Nakanishi, A. Recovery process of genetic diversity through seed and pollen immigration at the northernmost leading-edge population of *Fagus crenata*. Plant Species Biol. 36, 489–502 (2021).

107. Chybicki, I. J. et al. Disrupted connectivity within a metapopulation of a wind-pollinated declining conifer, Taxus baccata L. For. Ecosyst. 11, 100240 (2024).

108. Oddou-Muratorio, S. et al. Comparison of direct and indirect genetic methods for estimating seed and pollen dispersal in *Fagus sylvatica* and *Fagus crenata*. For. Ecol. Manag. 259, 2151–2159 (2010).

109. Toczydlowski, R. H. & Waller, D. M. Drift happens: Molecular genetic diversity and differentiation among populations of jewelweed (*Impatiens capensis* Meerb.) reflect fragmentation of floodplain forests. Mol. Ecol. 28, 2459–2475 (2019).

110. Naaf, T. et al. Sensitivity to habitat fragmentation across European landscapes in three temperate forest herbs. Landsc. Ecol. 36, 2831–2848 (2021).

111. Yıldız, B., Megens, H., Hvilsom, C. & Bosse, M. Genomic consequences of a century of inbreeding and isolation in the Danish wild boar population. Evol. Appl. 15, 954–966 (2022).

112. Damschen, E. I. et al. How fragmentation and corridors affect wind dynamics and seed dispersal in open habitats. Proc. Natl. Acad. Sci. 111, 3484–3489 (2014).

113. Rosot, M. A. D. et al. Riparian forest corridors: A prioritization analysis to the Landscape Sample Units of the Brazilian National Forest Inventory. Ecol. Indic. 93, 501–511 (2018).

114. Hardy, O. J. et al. Fine-scale genetic structure and gene dispersal inferences in 10 Neotropical tree species. Mol. Ecol. 15, 559–571 (2006).

115. Dick, C. W., Hardy, O. J., Jones, F. A. & Petit, R. J. Spatial Scales of Pollen and Seed-Mediated Gene Flow in Tropical Rain Forest Trees. Trop. Plant Biol. 1, 20–33 (2008).

116. Gelmi-Candusso, T. A., Heymann, E. W. & Heer, K. Effects of zoochory on the spatial genetic structure of plant populations. Mol. Ecol. 26, 5896–5910 (2017).

117. Gamba, D. & Muchhala, N. Pollinator type strongly impacts gene flow within and among plant populations for six Neotropical species. Ecology 104, e3845 (2023).

118. Browne, L., Ottewell, K. & Karubian, J. Short-term genetic consequences of habitat loss and fragmentation for the neotropical palm *Oenocarpus bataua*. Heredity 115, 389–395 (2015).

119. Solís-Hernández, W. & Fuchs, E.-J. Effective gene flow patterns across a fragmented landscape in southern Costa Rica for *Symphonia globulifera* (Clusiaceae); a species with mobile seed and pollen dispersers. Rev. Biol. Trop. 67, S95–S111 (2019).

120. Braun, M., Dantas, L., Esposito, T. & Pedrosa-Harand, A. Strong genetic differentiation on a small geographic scale in the Neotropical rainforest understory tree *Paypayrola blanchetiana* (Violaceae). Tree Genet. Genomes 16, 85 (2020).

121. González, A. V., Gómez-Silva, V., Ramírez, M. J. & Fontúrbel, F. E. Meta-analysis of the differential effects of habitat fragmentation and degradation on plant genetic diversity. Conserv. Biol. 34, 711–720 (2020).

122. Ony, M. A. et al. Habitat fragmentation influences genetic diversity and differentiation: Fine-scale population structure of *Cercis canadensis* (eastern redbud). Ecol. Evol. 10, 3655–3670 (2020).

123. Nazareno, A. G. & Jump, A. S. Species–genetic diversity correlations in habitat fragmentation can be biased by small sample sizes. Mol. Ecol. 21, 2847–2849 (2012).

124. Hoban, S. et al. Genetic diversity goals and targets have improved, but remain insufficient for clear implementation of the post-2020 global biodiversity framework. Conserv. Genet. 24, 181–191 (2023).

125. Lauterjung, M. B. et al. Temporal changes in population genetics of six threatened Brazilian plant species in a fragmented landscape. For. Ecol. Manag. 435, 144–150 (2019).

126. Hernández, M. et al. Population structure and genetic diversity of *Magnolia cubensis* subsp. *acunae* (Magnoliaceae): effects of habitat fragmentation and implications for conservation. Oryx 54, 451–459 (2020).

127. Mohana Kumara, P., Dayanandan, S., Vasudeva, R., Ravikanth, G. & Uma Shaanker, R. Population Genetic Diversity of *Dysoxylum Binectariferum*, an Economically Important Tree Species of the Western Ghats, India. in Molecular Genetics and Genomics Tools in Biodiversity Conservation (eds Kumar, A., Choudhury, B., Dayanandan, S. & Khan, M. L.) 251–266 (Springer Nature Singapore, Singapore, 2022).

128. Kubota, T. Y. K. et al. Pollen dispersal and mating patterns determine resilience for a large-yet-fragmented population of *Cariniana estrellensis*. Conserv. Genet. 25, 117–132 (2024).

129. Goetze, M., Capra, F., Büttow, M. V., Zanella, C. M. & Bered, F. High genetic diversity and demographic stability in *Aechmea kertesziae* (Bromeliaceae), a species of sandy coastal plains (restinga habitat) in southern Brazil. Bot. J. Linn. Soc. 186, 374–388 (2018).

130. Bard, N. W., Miller, C. S. & Bruederle, L. P. High genomic diversity maintained by populations of *Carex scirpoidea subsp. convoluta*, a paraphyletic Great Lakes ecotype. Conserv. Genet. 22, 169–185 (2021).

131. De Kort, H. et al. Life history, climate and biogeography interactively affect worldwide genetic diversity of plant and animal populations. Nat. Commun. 12, 516 (2021).

132. Wright, S. I., Kalisz, S. & Slotte, T. Evolutionary consequences of self-fertilization in plants. Proc. R. Soc. B Biol. Sci. 280, 20130133 (2013).

133. Barrett, S. C. H. & Harder, L. D. The Ecology of Mating and Its Evolutionary Consequences in Seed Plants. Annu. Rev. Ecol. Evol. Syst. 48, 135–157 (2017).

134. Silva, D. M. et al. Sexual and reproductive systems of woody species in *vereda* are distributed according to the life form and habitat occurrence. Austral Ecol. 47, 1528–1543 (2022).

135. Eckert, C. G. et al. Plant mating systems in a changing world. Trends Ecol. Evol. 25, 35–43 (2010).

136. Bittencourt, J. V. M. & Sebbenn, A. M. Patterns of pollen and seed dispersal in a small, fragmented population of the wind-pollinated tree *Araucaria angustifolia* in southern Brazil. Heredity 99, 580–591 (2007).

137. Ramos, S. L. F. et al. Paternity analysis, pollen flow, and spatial genetic structure of a natural population of *Euterpe precatoria* in the Brazilian Amazon. Ecol. Evol. 8, 11143–11157 (2018).

138. De Oliveira, S. S., Campos, T., Sebbenn, A. M. & d’Oliveira, M. V. N. Using spatial genetic structure of a population of *Swietenia macrophylla* King to integrate genetic diversity into management strategies in Southwestern Amazon. For. Ecol. Manag. 464, 118040 (2020).

139. Wang, X., Duan, F., Zhang, H., Han, H. & Gan, X. Fine-scale spatial genetic structure of the endangered plant *Tetracentron sinense* Oliv.(Trochodendraceae) in Leigong Mountain. Glob. Ecol. Conserv. 41, e02382 (2023).

140. Franklin, I. R., Soulé, M. E. & Wilcox, B. A. Conservation biology: an evolutionary-ecological perspective. Sunderland MA Sinauer Assoc. (1980).

141. Frankham, R. Evaluation of proposed genetic goals and targets for the Convention on Biological Diversity. Conserv. Genet. 23, 865–870 (2022).

142. Wold, J. et al. Expanding the conservation genomics toolbox: Incorporating structural variants to enhance genomic studies for species of conservation concern. Mol. Ecol. 30, 5949–5965 (2021).

